# BCMA signaling from the Golgi apparatus in B cells

**DOI:** 10.1101/2025.09.10.675116

**Authors:** Benoît Manfroi, Zeineb Chemkhi, Laurie Baert, Tejaswini Kakunuri, Olivier Micheau, Roland Govers, Harald Wajant, Pascal Schneider, Nathalie Sturm, Bertrand Huard

## Abstract

B-cell maturation antigen (BCMA) targeting gained a rapid approval for multiple myeloma, and is currently investigated in B-cell lymphomas. Here, we report that the trans Golgi network (TGN) retained BCMA due to a motif in its transmembrane/cytoplasmic domain in diffuse large B-cell lymphomas (DLBCL). To achieve signaling, one of its ligands, a proliferation inducing ligand (APRIL), bound to surface heparan sulfate proteoglycans (HSPGs), got endocytosed by the calveolin pathway and trafficked to the TGN via the retrograde route. In the TGN, APRIL/BCMA interactions activated the NF-κB pathway. BCMA ubiquitination followed by proteosomal degradation mediated signal termination. Growth impairment of DLBCL xenografts in *APRIL*-deficient animals confirmed the *in vivo* activity of TGN BCMA. Our study constitutes the first description for a signal activity from the Golgi apparatus among the TNF receptor family.

## Introduction

The member of the TNF receptor superfamily (TNFRSF), the B-cell maturation antigen (BCMA, TNFRSF17) is ubiquitously expressed by plasma cells (PCs). Interaction with one of its ligand, a proliferation inducing ligand (APRIL), regulates PC homeostasis ^1^. In addition to BCMA, APRIL has a second signaling receptor from the TNFRSF, the transmembrane activator and CAML interactor (TACI, TNFRSF13b) ^2^. TACI is involved in the isotype switch process and T-cell independent humoral responses ^3,4,5,6^. These roles of APRIL in humoral immunity have recently been confirmed with the description of the first APRIL immunodeficient patient ^7^. Finally, APRIL is quite unique in the TNFSF, since it has heparan sulfate proteoglycans (HSPGs) as coreceptors ^8^.

APRIL/BCMA and APRIL/TACI interactions promote the development of multiple myeloma (MM), a tumor of PC origin ^9,10,11^. While TACI is expressed on a subset of MM, BCMA is ubiquitous, and targeting of the latter has gained treatment approval with either cytotoxic, bispecific antibodies or T cells transduced with chimeric antigen receptors (CAR) ^12–14^. This implies a cell surface expression, which may be negatively regulated by gamma secretase activity ^15^. Notably, most recent studies revealed a puzzling mix localization for BCMA at the cell surface and in the Golgi apparatus in healthy PCs and MM ^16,17^. Several studies showed that paracrine APRIL produced by infiltrating neutrophils and a subset of macrophages also promotes development of B-cell lymphomas including diffuse large B-cell lymphomas (DLBCL) used in this study ^18–23^. Two main types of DLBCL have been reported. The germinal center like DLBCL harbors a gene expression profile close to germinal center (GC) B cells, while the activated B cell (ABC) like DBCL harbors a gene expression profile closer to the plasmablast stage ^24^. Due to the high efficiency of BCMA targeting in MM, BCMA expression has been widely studied in the B-cell lineage. With surface flow cytometry, BCMA was not detected in two independent studies ^25,26^. We previously reported BCMA intracellular expression with a small series of DLBCL cell lines ^18^. Notably, intracellular BCMA was also reported in human plasmacytoid dendritic cells ^27^. Regarding signaling, recombinant APRIL needs to be cross-linked to stimulate efficiently BCMA ^28^. In physiological conditions, such cross-linking is provided by the co-receptor activity of heparan sulfate proteoglycans (HSPGs) ^29,30^. The present study describes how BCMA when residing in the Golgi apparatus of DLBCL is triggered by paracrine APRIL.

## Results

### Endogenous BCMA is expressed in the trans Golgi network of DLBCL

Analysis of DLBCL primary tumor samples from untreated patients revealed tumor cells bound APRIL^A88^, which represents the mature soluble form of processed APRIL (figure 1A and 1sup). This was observed in 5/5 patients with 3 samples from the GC subtype and 2 samples from the ABC subtype. These cells expressed surface HSPGs, but showed surprisingly no cell surface expression of TACI and BCMA signaling receptors. Staining after cell permeabilization revealed a common intracellular pool for BCMA, while HSPG and TACI expression levels did not change. Such pattern of APRIL (co)receptor expression, hereafter called phenotype 1, was also observed in 11/18 DLBCL cell lines including BJAB (GC), Karpas 422 (GC), OCI-Ly1 (GC), OCI-Ly2 (GC), OCI-Ly7 (GC), Pfeiffer (GC), RC-K8 from the activated B cell (ABC) subtype, RIVA (ABC), Su-DHL-4 (GC), Su-DHL-7 (GC) and U2932 (ABC) cell lines. With cell lines, we also observed a phenotype called 2 not binding APRIL (figure 1B and 1sup). This concerned 7/18 cell lines included B593 (GC), OCI-Ly6 (ABC), OCI-Ly8 (GC), OCI-Ly18 (GC), Su-DHL-8 (GC) and Su-DHL-10 (GC) cell lines. The difference between the two phenotypes was the absence of surface HSPG in phenotype 2. Absence of BCMA surface expression was not due to high γ-secretase activity, because pharmalogical inhibition with DAPT up to 100 µM was not able to induce BCMA surface expression, while it did in the L363 plasma-cell leukemia cell line (figure 1C). Soluble BCMA was also not detectable in the supernatant on BJAB cells as opposed to L363 cells (figure 1D). To demonstrate the HSPG dependency of the APRIL binding to DLBCL of phenotype 1, we performed competition with the low-molecular weight soluble HS heparin and treatment with heparinase digesting the HS chain. It completely abrogated APRIL binding to cells from phenotype 1 (figure 1E). Likewise, APRIL^H98^, a truncated soluble form lacking the HSPG-binding domain but still able to bind TACI/BCMA, failed to bind cells from phenotype 1. In cells from phenotype 1, BCMA colocalized with Golgin-245, a marker associated to the trans-Golgi network (TGN) (figure 1F). The rat monoclonal antibody Vicky-1 used for BCMA detection by flow cytometry and cytofluorescence was not reactive with formalin-fixed paraffin-embedded (FFPE) tissues. We then validated with transfected cells for the immunohistochemistry application a rabbit polyclonal anti-BCMA made against an intracellular epitope (figure 2supA). Immunohistofluorescence with this antibody confirmed that DLBCL cells express intracellular BCMA *in situ* (Figure 1G). In healthy B cells, BCMA expression profile revealed that transcription starts in GC B cells in both centrocytes and centroblasts and peaked in plasma cells (figure 2supB). Immunohistofluorescence showed also an intracellular localization in CD20^+^ GC B cells (figure 2supC). Flow cytometry confirmed that most CD20^+^ B cells expressed BCMA intracellularly (figure 2supD). The few CD20^+^ B cells harboring surface expression of BCMA had in fact a morphology reminiscent with the more differentiated stage plasmablast.

**Figure 1:**
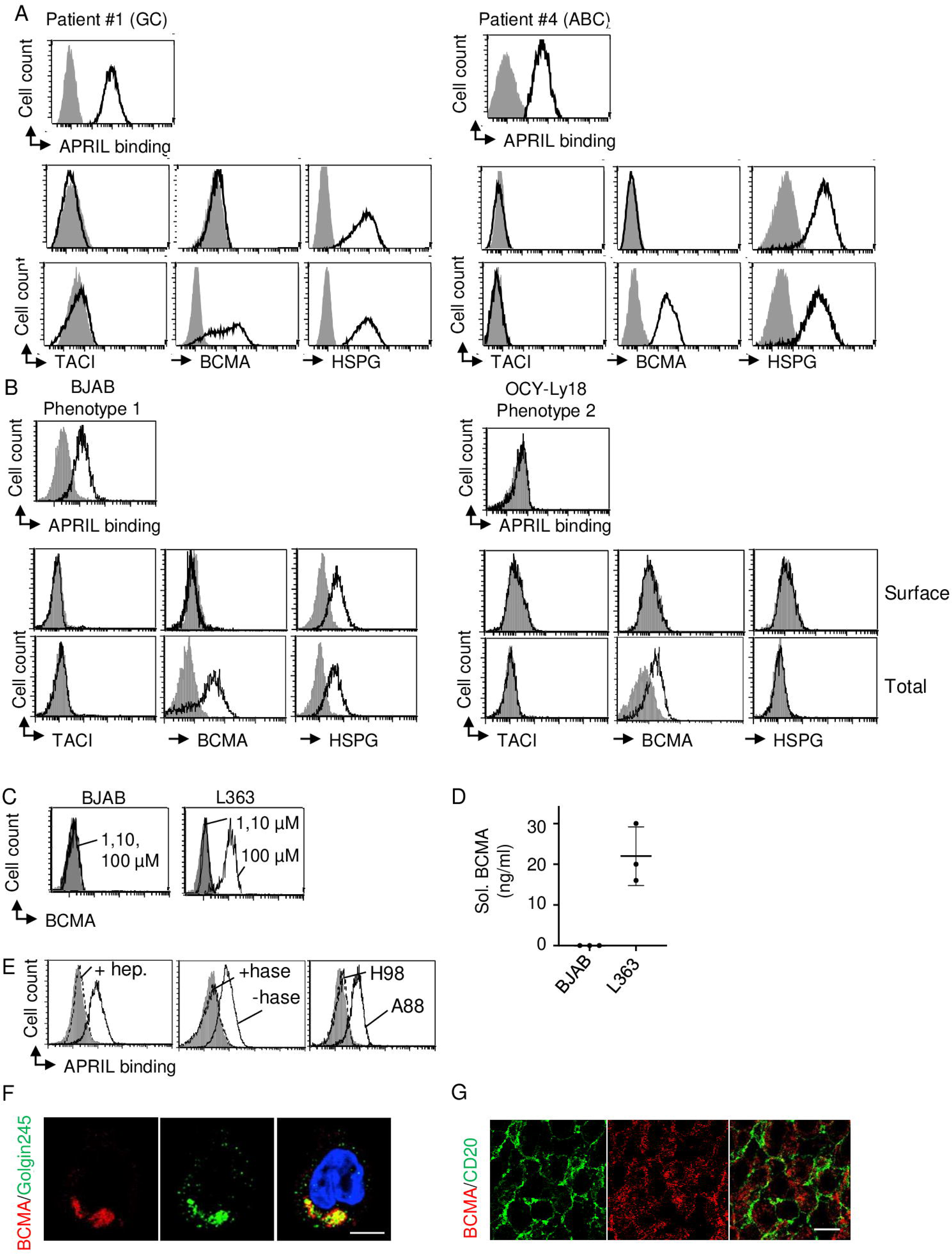
Endogenous BCMA is retained in the trans Golgi network of DLBCL. A) Primary samples from DLBCL patients were stained with recombinant soluble APRIL^A88^ (upper panel) and analyzed for the surface and total expression of the indicated receptors by flow cytometry (lower panel). B) DLBCL cell lines were analyzed as in A. C) The indicated cell lines were treated with the γ-secretase inhibitor DAPT at 1, 10 and 100 µM prior to surface BCMA staining and flow cytometry. Overlayed histograms are shown. D) Detection of soluble BCMA by ELISA with supernatants conditioned for 3 days. E) BJAB were stained with recombinant APRIL ^A88^ in the presence of heparin (hep.) (left panel), with or without prior heparinase (hase) treatment (middle panel), and with APRIL^A88^/APRIL^H98^ (right panel). F) Fixed/permeabilized BJAB cells were stained for BCMA and Golgin-245 and analyzed by confocal microscopy. Scale bar = 5µm. G) A DLBCL biopsy was costained for CD20 and BCMA and analyzed by confocal microscopy. Scale bar = 10 µm. Figures A//B/C/D/E/F are representative of three experiments. Figure G is representative of fifteen DLBCL patients.

### Transfected BCMA reproduces a Golgi-like pattern

Transfection of full-length YFP-tagged human BCMA in BJAB cells from phenotype 1 reproduced an intracellular perinuclear localization, reminiscent of the Golgi apparatus (figure 2A). In additional transfection experiments, flow cytometry with the anti-BCMA did not detect any surface expression of transfected full-length human BCMA, while it did when the transmembrane/cytoplasmic domain was replaced by a glycolipid anchor (figure 2B, upper left panel). In this experiment, transfected full-length human TACI, the other signaling APRIL receptor, localized at the cell surface (figure 2B, upper right panel). We also transfected full-length mouse BCMA in BJAB. Again, transfected mouse BCMA was not detected at the cell surface, while a strong signal was obtained after cell permeabilization (figure 2B bottom panel). Similar results were obtained with the two other DLBCL cell lines SU-DHL7 (GC) and RIVA (ABC) from phenotype 1 (figure 3supA). Altogether, this showed that BCMA is retained in the TGN by a motif present in its transmembrane/cytoplasmic domain when expressed in DLBCL cells. A similar intracellular staining is observed in a fraction of GC B cells.

**Figure 2:**
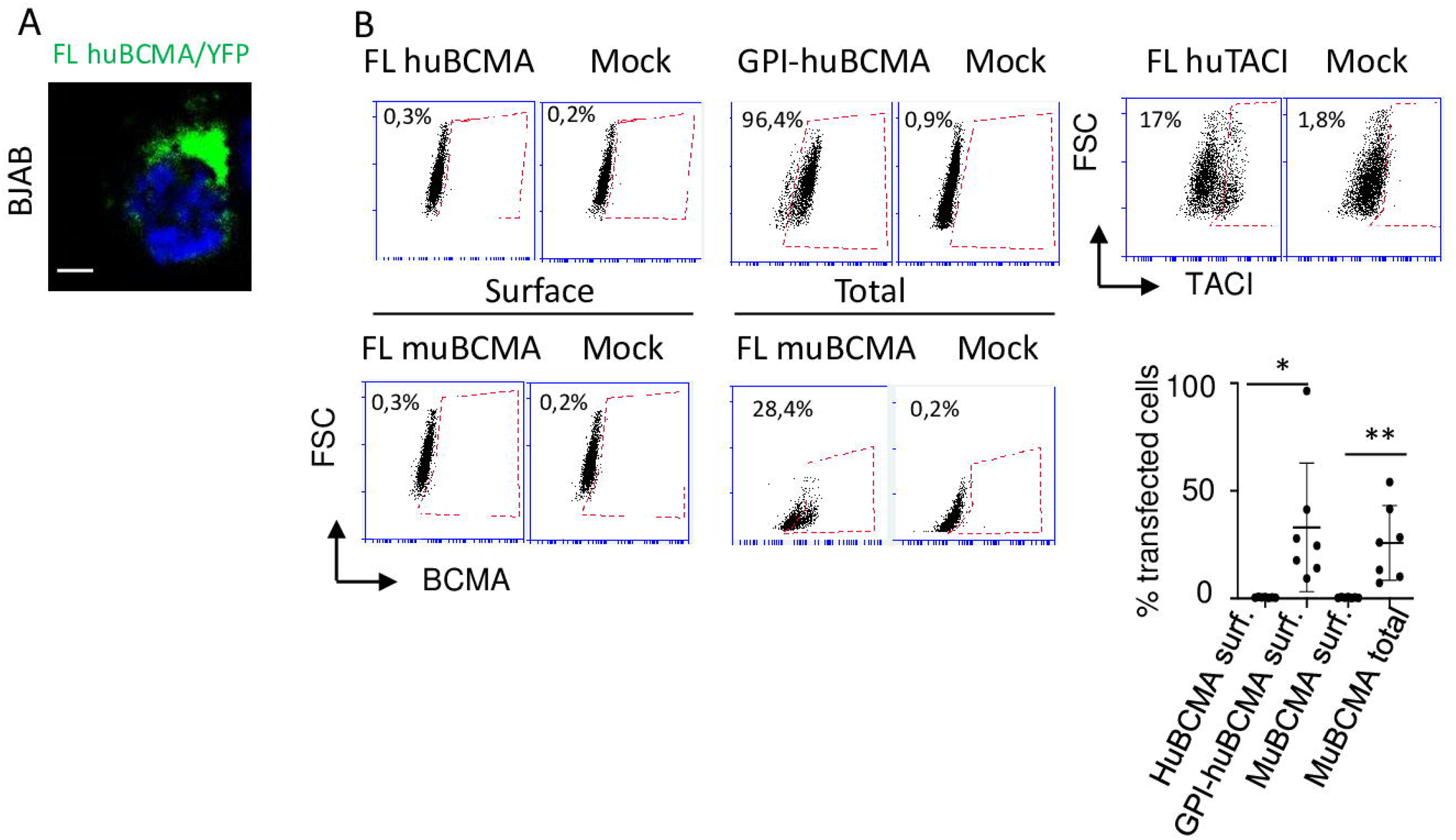
Transfected BCMA harbors a Golgi localization in DLBCL. A) BJAB cells were transfected with YFP-full-length BCMA and analyzed by confocal microscopy 48 hr later. B) Live (surface) or permeabilized (total) BJAB cells transfected with the indicated constructs were stained for huBCMA, huTACI or muBCMA, and analyzed by flow cytometry (upper and bottom left panels). Dot plots are shown. FL: full length, GPI: glycophosphatidyl inositol. % of transfection and localization of the BCMA molecules is reported from three experiments performed with BJAB, RIVA and SuDHL-7 cell lines (bottom right panel). An unpaired *t*-test was performed. Figures A is representative of three experiments.

### APRIL is internalized in DLBCL and interacts with BCMA in the TGN

We next studied the fate of surface bound APRIL^A88^ detected by its flag tag in BJAB cells. About half of BJAB cells lost membrane-bound APRIL when incubated for 45 min at 37°C (figure 3A). Permeabilization of these cells allowed full recovery of the APRIL signal. Inhibition of caveolin-mediated endocytosis with filipin III strongly impaired APRIL internalization, while inhibition of clathrin-mediated endocytosis with sucrose had no effect. After 15 min, confocal microscopy analysis confirmed APRIL internalization at 37°C, while APRIL stayed at the cell surface at 4°C, colocalizing with the surface marker cholera toxin B subunit (CTxB) (figure 3B). In this experiment, OCI-Ly18 cells belonging to phenotype 2 did not show any APRIL internalization at 37°C (figure sup 3B). Kinetic analysis revealed that APRIL first capped in lipid-rich, CTxB stained, raft regions after 5min, and then colocalized with calveolin-but very rarely with clathrin-coated vesicles after 10 min at 37°C (figure 3C, left panel). After 15 min, APRIL localized in EEA1^+^ early but not LAMP1^+^ l late endosomes. After 30 min of incubation, APRIL was mostly found in LAMP1 but more rarely in EEA1^+^ endosomes (figure 3C, right panel). Finally, we never found APRIL in Rab11^+^ recycling endosomes. After 45 min at 37°C, APRIL co-localized with the Golgin-245 marker, and intracellular BCMA in BJAB cells. This colocalization in the Golgi apparatus was also observed in SU-DHL7 and RIVA DLBCL cell lines (figure sup 3C). In these experiments, pretreatment of cells with Retro2, a specific inhibitor of the endosome to Golgi retrograde transport, favored LAMP1/APRIL over BCMA/APRIL colocalizations at the 45 min time point (figure 3D). APRIL internalization by BJAB cells was strictly dependent on HSPGs, since inhibited by heparin but not by a soluble form of TACI blocking APRIL binding to TACI/BCMA (figure sup 3D).

**Figure 3:**
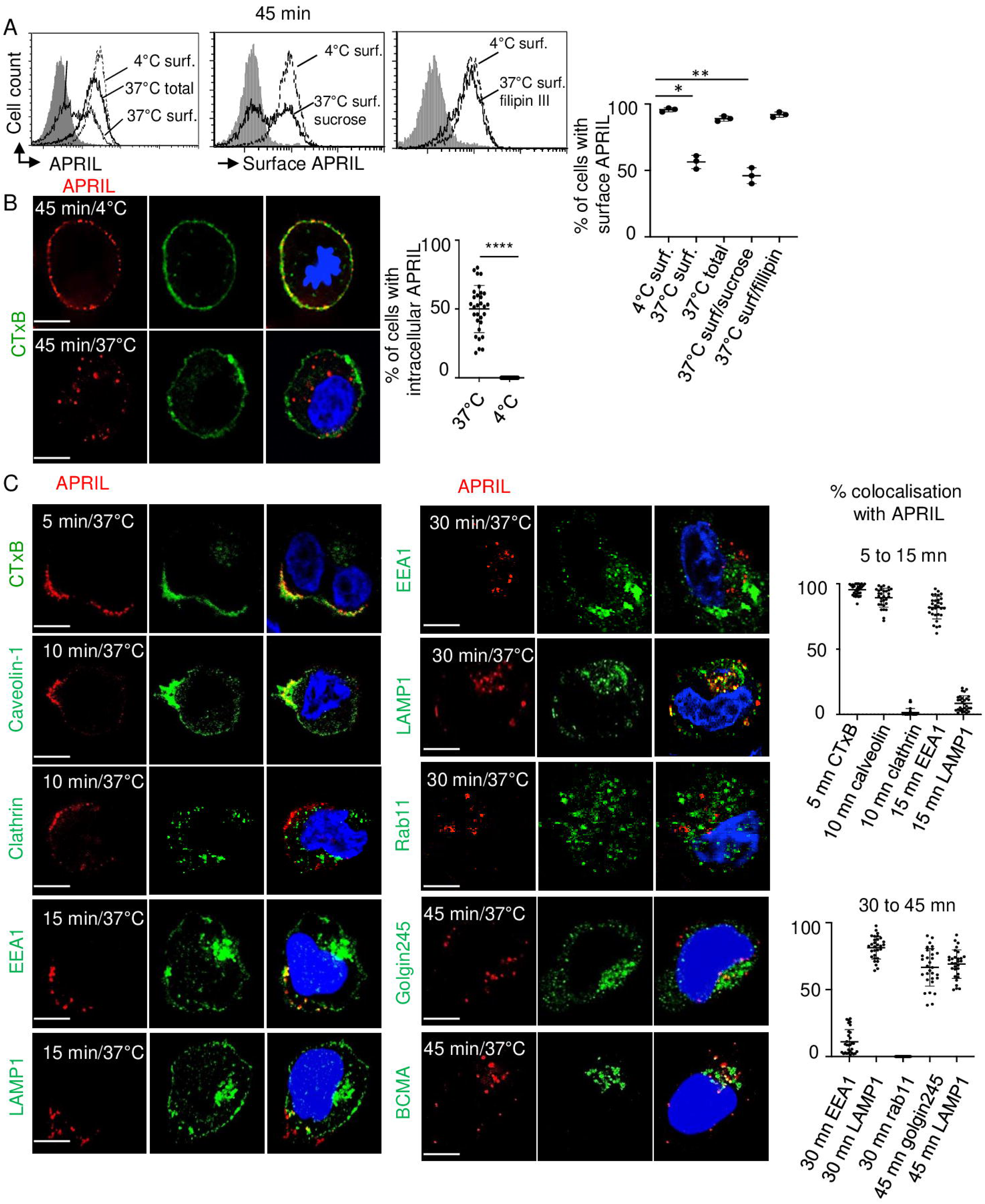
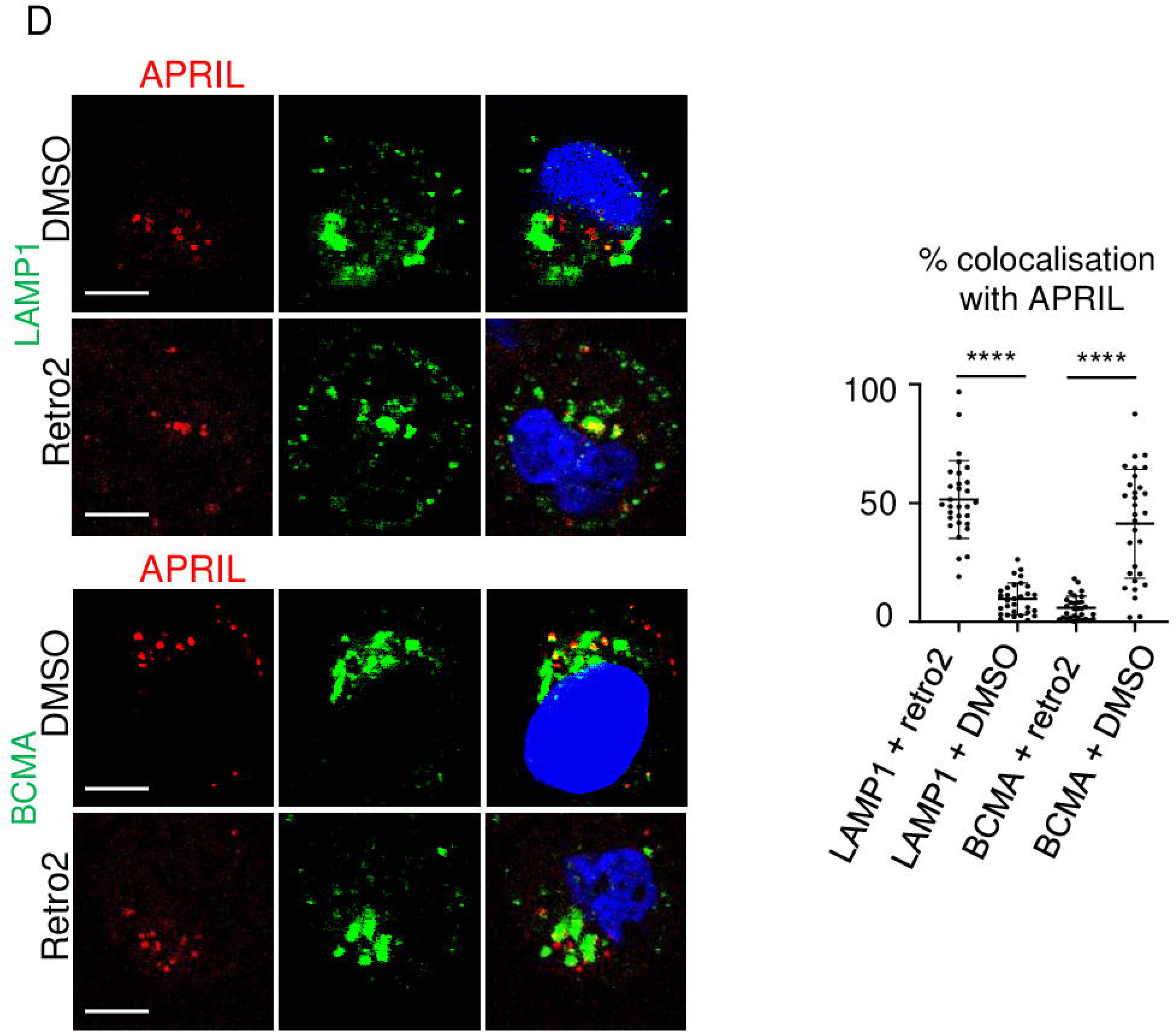
Internalized APRIL traffics to the trans-Golgi network via the retrograde route in DLBCL. APRIL^A88^ was added to BJAB cells at 4°C. Cells were washed and incubated as indicated. A) APRIL (anti-flag reactivity) on viable cells was assessed at the cell surface (surf.) and after cell permeabilization (total) by flow cytometry (left panel). BJAB cells pretreated with or without endocytosis inhibitors were stained for membrane-bound APRIL expression (middle and right panels). A representative experiment is shown (left panel). Individual values for three independent experiments are shown (right panel). B/C) APRIL ^A88^-incubated BJAB cells were stained for the indicated markers at the indicated times and temperature before analysis by cytofluorescence. Representative images are shown (left panels). The % of colocalization calculated from 30 cells is shown (right panels). D) Retro2- or vehicle-pretreated BJAB cells were stained for APRIL/LAMP-1 and APRIL/BCMA. Representative images are shown (left panel). The % of colocalization calculated from 30 cells is shown (right panel). Colocalizations were analyzed and quantified by cytofluorescence. Means +/- SD are shown. Scale bar = 5µm. A paired one-way Anova test was performed in A. A paired *t-*test was performed in B/D. Figures A/C/D are representative of 3 experiments. Figure B is representative of 5 experiments.

### HSPGs are responsible for APRIL internalization in DLBCL

Next, we investigated HSPG fate in APRIL-stimulated cells with an anti-HS mAb. In response to APRIL, about half of the cells also lost membrane HSPGs (figure 4A), and about half of the APRIL signal colocalized intracellularly with HSPG after 15 min at 37°C while none of the APRIL and HSPG signals were detected intracellularly at 4°C (figure 4B). Kinetics analysis revealed that after 5 min, HSPG also capped in lipid rich raft, CTxB stained, regions (figure 4C). After 15 min at 37°C, internalized HSPG also localized in EEA-1^+^ early endosomes. It was only later than 15 min that we observed an HSPG trafficking different from APRIL. Although, both left early endosomes, HSPGs barely localized in LAMP1^+^ late endosomes. Instead, we observed a significant fraction of HSPGs in Rab11^+^ recycling endosomes after 30 min. This process was never detected spontaneously in the absence of APRIL. In fact, APRIL fully disappeared from the cells surface after 180 min at 37°C, while HSPG cell surface expression recovered with time (figure 4D). We did not detect loss of surface bound APRIL, if unbound APRIL was not removed by a washing step prior to the 37°C incubation (figure sup 3E). This constituted another argument for a cell surface recycling of HSPG. A cell-feeding experiment with the anti-HSPG 10e4 unable to block APRIL binding to its coreceptor confirmed the cell surface recycling process for HSPG. In the beginning of the experiment, all cells presented the anti-HSPG at their surface in the absence of APRIL (figure 4E). A first incubation for 30 min at 37°C with APRIL led to some internalization of the anti-HSPG. An acid treatment removed most of the remaining surface anti-HSPG. A second incubation at 37°C for 60 min after this acid treatment allowed a redetection of the anti-HSPG at the surface of cells. In BJAB cells, the HSPG proteic core was CD44v3, an alternative spliced form of CD44 known to selectively harbor HS chains ^31^ (figure 5A). Notably, we observed an identical endocytosis pathway following APRIL stimulation for the proteic core of CD44v3 compared to the HS chain with a sequential colocalization in EEA1^+^ early and Rab11^+^ recycling but barely in LAMP1^+^ late endosomes (figure 5B). Taken together, this shows that HSPGs mediate APRIL internalization until the early endosome stage. After early endosomes, the APRIL/HSPG complex dissociates. HSPGs recycle to the cell surface, while APRIL traffics to the TGN to meet BCMA. Such APRIL internalization observed *in vitro* is fully consistent with the intracellular accumulation of APRIL previously reported *in situ* in DLBCL ^18^.

**Figure 4:**
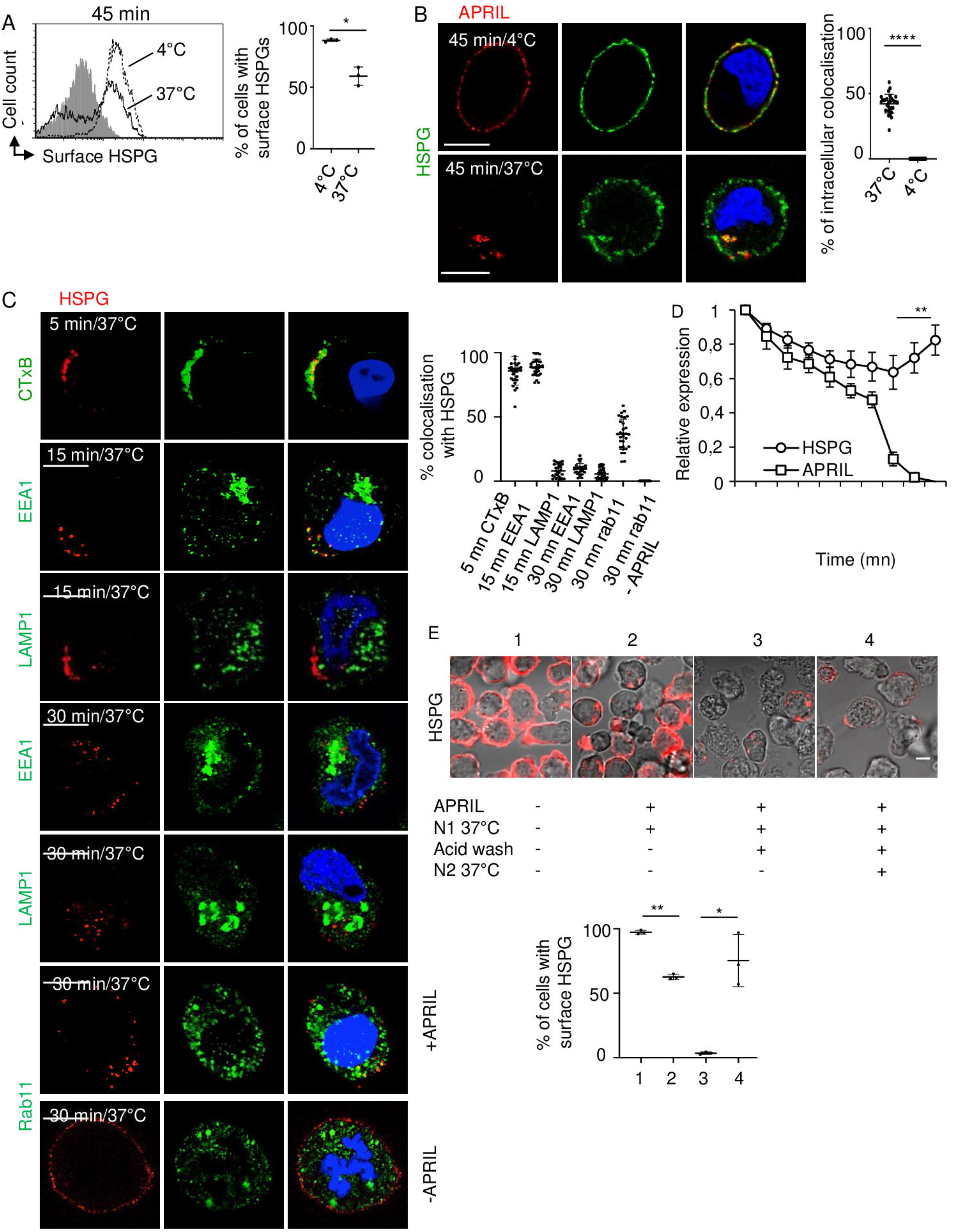
HSPGs internalize APRIL and recycle to the cell surface. APRIL^A88^ was added to BJAB cells at 4°C. Cells were washed and incubated as indicated. A) HSPG on viable cells was assessed by flow cytometry (left panel). Three independent experiments are shown (right panel). B) APRIL ^A88^-incubated BJAB cells were stained for APRIL and HSPGs at the indicated times and temperature before analysis by cytofluorescence. Representative images are shown (left panel). The % of the HSPG signal colocalizing intracellularly with APRIL was calculated from 30 cells (right panel). C) APRIL ^A88^-incubated BJAB cells were stained for HSPGs and the indicated markers at the indicated times and temperature before analysis by cytofluorescence. Representative images are shown (left panel). The experiment for HSPG/Rab11 costaining was performed in the presence or absence of APRIL ^A88^ stimulation. The % of colocalization calculated from 30 cells is shown (right panel). D) Kinetics of relative surface HSPG and APRIL expression after APRIL stimulation at 37°C. The 1 value corresponds to staining when cells were kept at 4°C. E) BJAB cells were fed with an anti-HSPG (IgM) and HSPG expression was analyzed with a fluorochrome-conjugated anti-IgM in the indicated conditions by cytofluorescence. Representative images are shown (left panel). The % of cells with surface HSPG was calculated from 3 experiments (right panel). Paired *t*-tests were performed. Scale bars = 5µm. All pictures are representative of at least 3 independent experiments except D representative of two independent experiments performed in triplicates. Mean ± SD is shown.

**Figure 5:**
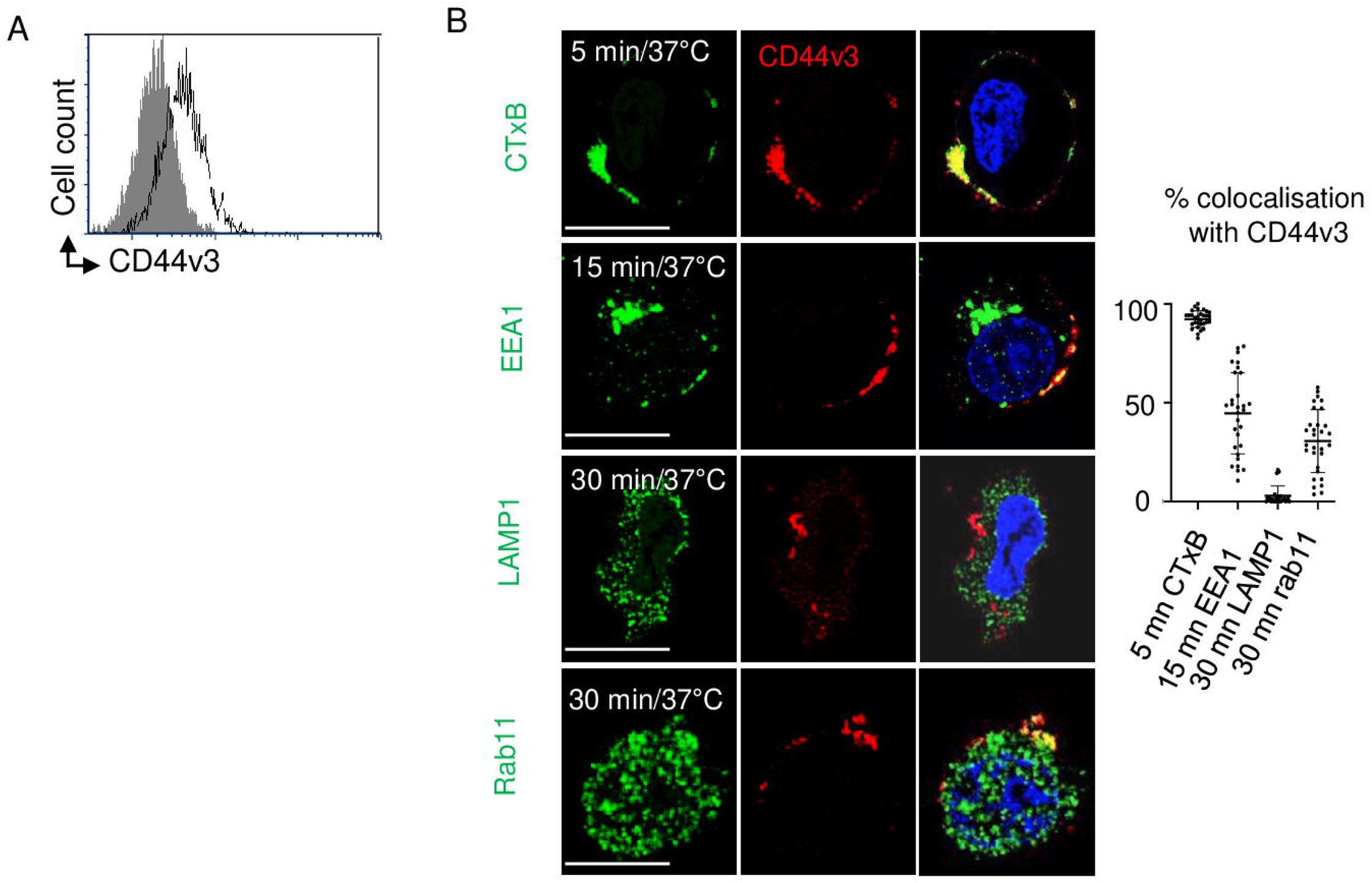
The protein core of HSPG CD44v3 is also internalized. A) CD44v3 surface reactivity of BJAB cells was analyzed by flow cytometry. B) BJAB cells were incubated at 37°C for the indicated times with soluble APRIL^A88^, costained for the indicated markers and analyzed by cytofluorescence. Representative images are shown (left panel). The % of colocalisation calculated from 30 cells is shown (right panel). Scale bar = 5µm. Figures are representative of two independent experiments.

### APRIL/BCMA interactions in the TGN activate the NF-κB pathway

We next evaluated the outcome of APRIL localization in the TGN. Analysis of APRIL immunoprecipitates showed that APRIL/BCMA interactions started to be detectable after 30 min at 37°C (figure 6A). Out of the 6 TNF receptor-associated factors (TRAFs), we found TRAF1 and 2 present in the complex, consistent with previous reports on BCMA signaling ^20,32,33^. We also detected in the complex the cellular inhibitor of apoptosis protein 1 (cIAP1), another component frequently involved in the signaling of TNFRSF receptors ^34^. Timing of APRIL/BCMA interactions correlated with activation of NF-κB as detected by induction of phospho-IκB-α and loss of total IκB-α. In this experiment, co-immunoprecipitated BCMA was identified as a ladder with bands showing molecular weight gain of up to 25 kDa, reminiscent of ubiquitination. Pretreatment of BJAB cells with the proteasome inhibitor MG132 blocked loss of BCMA staining observed in APRIL-treated cells incubated at 37°C (figure 6B). Thus, proteosomal degradation following polyubiquitination of BCMA may contribute to the termination of the APRIL-induced BCMA signaling. We next analyzed qualitatively APRIL/BCMA signaling properties in BJAB cells. By studying cytokine secretion upon APRIL stimulation, we never detected upregulation of the proinflammatory cytokines IL-1β, IL-6, IL-8, IL-12 and TNF, but reproducibly observed upregulation of IL-10 in phenotype 1 with BJAB cells but not in phenotype 2 with OCI-Ly18 cells (figure 6C). IL-10 upregulation was NF-κB-dependent, since inhibited with parthenolide, an antagonist of this pathway. We also used transfection with an undegradable mutant of IkB in BJAB cells to further assess NFkB dependency. For some reasons, the flag tag present in this mutant was not detectable by flow cytometry. We cloned the transfected line and assessed flag reactivity in clone lysates by denaturating Western blot (figure 6D, left panel). A band at the expected size of 35 kDa was detected by the anti-flag antibody. Two clones expressing the mutant IkB showed loss of IL-10 upregulation upon APRIL stimulation compared to a control GFP-transfected clone, thus confirming that the APRIL signaling relies on the NFkB pathway (figure 6D, right panel). We further observed a positive correlation between *APRIL* and *IL-10* mRNA level *in situ* obtained with FACS-sorted CD20^-^ microenvironment and CD20^+^ tumor cells, respectively, from DLBCL biopsies (figure 6E). To test the function of IL-10 produced by APRIL-stimulated DLBCL, we performed a mixed lymphocyte reaction (MLR) with BJAB cells and allogeneic T cells. In the presence of APRIL-treated BJAB cells, the proliferative percentage of CD4^+^ T cells decreased, and addition of an anti-IL-10 antagonist allowed to partially recover T-cell proliferation (figure 6F). In this MLR experiment, we also detected a significant decrease in secretion of the T-cell-derived cytokines IL-6, IL-2, IL-17 and IFNγ in the MLR supernatant, when BJAB cells were stimulated with APRIL prior to the MLR. IL-10 antagonism also partially allowed cytokine secretion recovery. Altogether our data shows that the intracellular APRIL/BCMA signaling pathway is functional at least *in vitro*, and increases DLBCL immunosuppressive properties.

**Figure 6:**
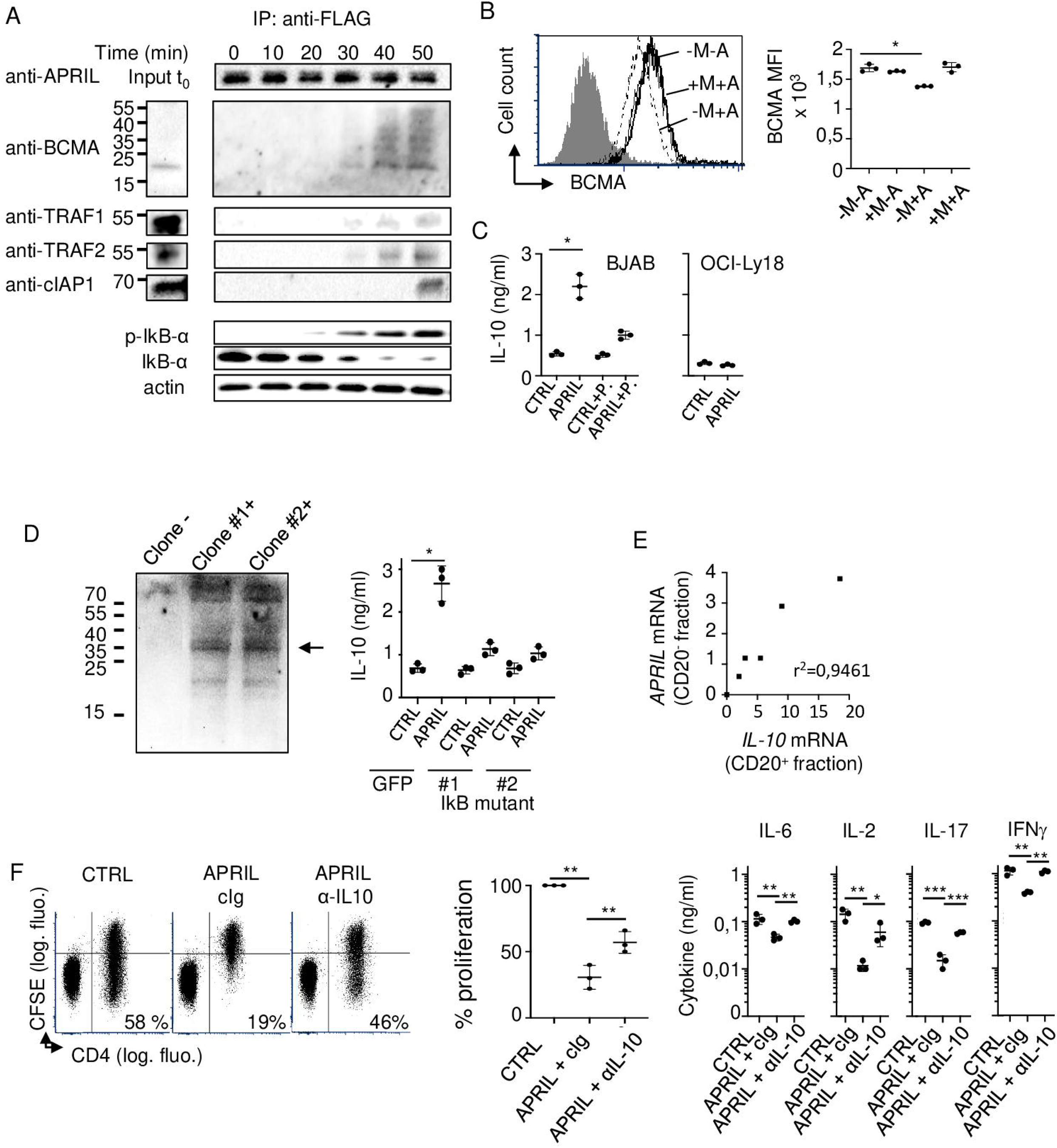
Intracellular APRIL/BCMA interactions activates the NF-κB signaling pathway. BJAB cells were incubated with APRIL^A88^ at 37°C. A) APRIL was immunoprecipitated at different time points and analyzed by Western blot with the indicated antibodies. Inputs at t_0_ are shown (upper panel). Phosphorylated and total Iκb-α in whole cell extracts are also shown with actin as a control (lower panel). B) MG132 (M)-pretreated BJAB cells were incubated with APRIL^A88^ (A) for 30 min and BCMA total expression after cell permeabilization was assessed by flow cytometry. A representative overlay histogram plot is shown (upper panel). Fluorescence intensity (MFI) is shown (right panel). C) APRIL ^A88^ - or CTRL-stimulated BJAB (left) and OCI-Ly18 (right) cells were assessed for IL-10 secretion by ELISA after 48 hr. BJAB cells were also tested in the presence or absence of parthenolide (P.). D) Western-blot analysis of lysates of indicated cells is shown (left panel). APRIL ^A88^ - or CTRL-stimulated BJAB cells transfected with GFP or the Ikb mutant were assessed for IL-10 secretion by ELISA after 48 hr (right panel). E) *APRIL* from the microenvironment (CD20^-^ fraction) and *IL-10* from the tumor (CD20^+^) mRNAs were quantified by qRT-PCR. Relative *APRIL* and *IL-10* mRNA expressions in primary samples from DLBCL patients is shown. Results are representative of three experiments. F) A MLR was performed between BJAB cells and allogeneic PBMCs. CD4^+^ T-cell proliferation was assessed with CFSE dilution by cytofluorimetry after 72 hr. Representative dot plots are shown (left panel). The % +/- SD of dividing CD4^+^ T cells is also shown (middle panel). 100% represents the response obtained in the presence of CTRL-treated BJAB cells. Concentrations of the indicated cytokine in the MLR supernatant is also shown (right panel). A one-way Anova tested was performed in B. Paired *t* tests were performed in C and D. Pearson correlation was performed in E. Dunnett’s multiple comparison test was performed in F. Mean +/- SD of three independent experiments is shown in B, C, D, and F. Uncropped Western blots are shown in figure 4sup.

### APRIL/BCMA interaction in the TGN promotes DLBCL development

We next assessed the APRIL-dependency of DLBCL from phenotype 1 *in vivo*. For that purpose, we backcrossed *APRIL* KO mice onto the immunodeficient background NOD/SCID. Interaction of mouse APRIL with human BCMA has already been reported by flow cytometry ^35^. We further observed that this interaction of mouse and human APRIL to huBCMA is equally efficient in ELISA, and equally able to induce death of Jurkat T cells transfected with a fusion protein comprising the extracellular domain of human BCMA and the intracellular domain of Fas (JKT-huBCMA/Fas) (figure 5sup). BJAB cells did not engraft in these immunodeficient mice, but the OCI-Ly7 (GC) cell line also from phenotype 1 engrafted efficiently. Growth of OCI-Ly7 was significantly impaired in the absence of mouse APRIL since only 2 and all mice harbored a palpable tumor reaching more than 0,5 cm^2^ in size in *APRIL*-deficient and WT mice, respectively (figure 7). Taken together, this shows that intracellular APRIL signaling promotes DLBCL growth.

**Figure 7:**
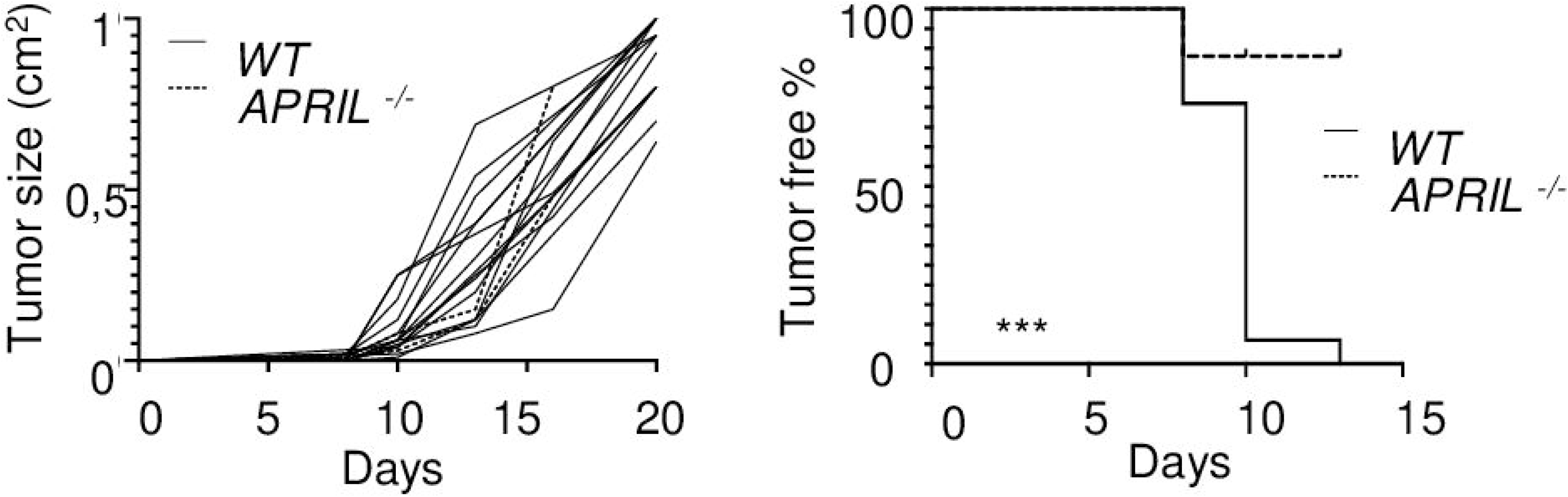
Intracellular APRIL signaling promotes DLBCL growth. OCI-Ly7 cells were injected subcutaneously in the indicated NOD/SCID mice. Tumor size at the indicated time is shown for 15 mice (left panel). Tumor engrafment is also shown (right panel). A Log-rank test was performed.

## Discussion

A recent review indicated that localization of BCMA in B-cell lymphomas is contradictory with either the description of surface or intracellular localizations ^36^. Here, we are showing an exclusive intracellular localization, more specifically in the TGN, for BCMA in DLBCLs. This confirms the initial report made upon cloning of the BCMA gene and showing a Golgi localization in mature B cells. B-cell lymphomas are currently test for anti-BCMA therapies. Burkitt, mantle cell lymphomas and chronic lymphocytic leukemia gave encouraging preclinical results ^37,38^. Results regarding DLBCL remains limited. Nevertheless, the CAR T-cell activation with IFNγ secretion reported upon coculture with the use of an affinity improved CAR BCMA indicates that a pool of BCMA should exist at the DLBCL cell surface, either from a stable or recycling pool ^39^. This pool might not be detectable reproducibly with conventional antibody-based staining.

Early work with the TGN38 glycoprotein indicated that residues in the transmembrane domains and cytoplasmic tail act independently to mediate Golgi retention ^40^. If one considered a global Golgi localization, several pathways have been reported since then without reaching any consensus regarding primary sequences ^41,42^. Golgi resident proteins are usually type II proteins, while BCMA is a type III protein. If one challenge further the human and mouse BCMA sequences to Golgi localization motifs, BCMA does not have a so-called GRIP domain in its luminal part. It has neither a glutamine in its transmembrane domain nor a dibasic motif in its membrane proximal cytoplasmic tail. The exoplasmic part of its transmembrane domain is composed of residues rather not bulky. Regarding length of domains potentially involved in Golgi retention, the transmembrane domain and cytoplasmic tail are longer than the average of Golgi retained proteins. Altogether, this indicates that BCMA may possess a specific Golgi retention pathway, warranting further investigations. We are further showing that TGN BCMA is signaling active. Such intracellular signaling is rendered possible by the internalization ability of HSPGs known as coreceptors of one of the BCMA ligand, APRIL, highly consistent with their well-described endocytic property ^43^. This property has also been demonstrated for CD44, whose variant 3 bears HS chains and is expressed in B cells ^44^. Once dissociated from HSPGs, APRIL uses the retrograde transport route from endosomes to the Golgi apparatus, a transport route initially described for secreted bacterial toxins to reach the host cell cytosol ^45^. Notably, Golgin-245 itself participates in the retrograde transport route ^46^. Hence, once dissociated from HSPG, internalized APRIL may follow the Golgin-245 pathway to interact with BCMA in the TGN. We also observed signs of BCMA ubiquitination with the appearance of higher molecular weight band. Weight gain for BCMA is quite important, up to 25 kDa, reminiscent of a polyubiquitination rather than mono-ubiquitination state. Associated TRAF1 does not possess in its N-terminus part a RING finger domain containing the core of the ubiquitin ligase activity. However, associated TRAF2 and cIAP1 possess this activity ^47^. Hence, BCMA polyubiquitination may serve to terminate the signal by sending it to proteasomal degradation. TNFRs are usually not ubiquitinated ^47^. It is rather the associated TRAFs that are targeted. However, the high number (n=9) of lysine, site of ubiquitination on proteins ^48^, present in the BCMA cytoplasmic tail makes it a potential target molecule for this process. Here, we are reporting NF-κB activation and IL-10-based immunosuppression by TGN BCMA in APRIL-stimulated DLBCL. The production of IL-10 by DLBCL observed here is quite consistent with previous reports showing IL-10 upregulation in DLBCL patients and correlation to poor prognosis, also explained by the existence of an autocrine proliferation loop for IL-10R^+^ DLBCL tumor cells ^49,50,51^.

Protein retention in the Golgi apparatus is obviously needed for biosynthetic enzymes involved in glycan processing occurring at that specific place ^41^. However, signaling from the Golgi apparatus is now an emerging concept for receptors previously thought to act at the cell surface. This is the case for G protein-coupled receptors, the largest family of signaling receptors, with some members described as resident in the Golgi apparatus ^52^. For the latter, spatial modulation of cell signaling has been proposed. This might not be the case for BCMA, since APRIL stimulation of MM expressing surface BCMA looks highly similar to APRIL stimulation of DLBCL expressing Golgi BCMA with NF-κB activation and IL-10 upregulation ^10^. DLBCL arise from the transformation of healthy GC B cells. Notably, an intracellular localization of BCMA in healthy GC B cells is also observed, indicating that intracellular retention of BCMA is not acquired after cell transformation. Further investigations are warranted to understand why along the B-cell differentiation process during a humoral response, cells form the GC stage harbor first Golgi expression of BCMA before showing a surface expression of the latter at the latest stage of differentiation, the PC. In total, our study shows that signaling from the Golgi apparatus also occurs among the TNFRSF.

## Materials and methods

### Antibodies and reagents

Origin and use of antibodies against CD20 (clone L26, mouse IgG2a) and HSPG (clone 10e4, mouse IgM) have been described elsewhere ^18,53^. Validation of the TACI (clone C4, mouse IgG1) and BCMA (clone Vicky-1, rat IgG1) antibodies on transfected cells for flow cytometry was previously described ^18^. TACI-Ig used at 50 µg/ml was from Adipogen. Anti-CD44v3 was from Calbiochem. FITC-labeled cholera toxin B subunit (CTxB, Sigma) was used at 10µg/ml. Anti-caveolin-1 (N-20, rabbit IgG, Santa Cruz Biotechnology), anti-clathrin (goat IgG, Sigma), anti-EEA1 (rabbit IgG, Cell Signaling Technology), anti-Golgin-245 (C-13, rabbit IgG, Santa Cruz Biotechnology), anti-LAMP1 (N-19, goat IgG, Santa Cruz Biotechnology) and anti-rab11 (C-19, rabbit IgG, Santa Cruz Biotechnology) were used at 1µg/ml in immunofluorescence staining. The anti-TRAF1 (clone 45D3, rabbit IgG), -TRAF2 (clone C192, rabbit IgG), -TRAF3 (rabbit IgG), -TRAF4 (clone D1N3A, rabbit IgG), - TRAF5 (clone D3E2R, rabbit IgG), -TRAF6 (clone D21G3, rabbit IgG), anti-phospho-IκB-α (Ser32, clone 14D4, rabbit IgG), anti-IκB-α (clone 44D4, rabbit IgG) and anti-actin (clone 13E5, rabbit IgG) were from Cell Signaling Technology. Anti-cIAP1 and cIAP2 (both goat IgG) were from R&D systems. All antibodies in Western blot analysis were used at 0.2µg/ml. Neutralizing anti-IL-10 (clone 25209, mouse IgG2b) was from R&D systems and used at 5µg/ml. The rabbit polyclonal anti-BCMA (Abcam) is directed against an intracellular epitope, and was used at 3 µg/ml. Conjugated secondary reagents were from ThermoFisher. Fluorochrome conjugated anti-CD20 antibody was from BD biosciences. Anti-muBCMA (Clone Vicky-2, rat mAb) was from Enzo Life Sciences. For endocytosis analysis, cells were treated with Filipin III (2µg/ml in OptiMEM, Sigma) or with a hypertonic solution (300mM sucrose in OptiMEM). Flag-tagged recombinant trimeric human APRIL was transiently produced in HEK 293T cells with serum-free OptiMEM medium by using polyethylenimine-based transfection. APRIL^A88^ (88-233), which represents the mature form of APRIL, was used in this study. APRIL^H98^ (98-233) is a mutant form deleted for the HSPG-binding domain. Flag-tagged EDA (245-391) was used as a control ^35^. Crude supernatant was concentrated 10X with Amicon Ultra-15 (30 kDa cut-off, Millipore) before binding/internalization studies. Binding of Flag-tagged recombinant APRIL was detected with a biotinylated anti-Flag antibody (clone M2, mouse IgG1, Sigma) followed by fluorochrome-conjugated streptavidin (BD biosciences). For functional analysis, recombinant APRIL was purified on anti-Flag agarose M2 affinity gel (Sigma) using standard procedures. The γ-secretase inhibitor DAPT (Sigma) was incubated with cells for 4 hr at 37°C prior to flow cytometry staining. The IL-10 and soluble BCMA ELISA kits were from Bio-techne.

### Cells

All DLBCL cell lines used in this study were described elsewhere ^54^. They were not passaged more than 10 times upon receipt. The plasma-cell leukemia L363 was obtained from the American Tissue Culture Collection. Prior to investigation, they were checked for absence of mycoplasma contamination with the plasmotest kit (InvivoGen). The NF-κB pathway inhibitor parthenolide (Sigma) was used at 5 µM. MG132 proteasome inhibitor (Calbiochem) was used at 10µM for 2 h prior to APRIL stimulation. Retro-2, a specific inhibitor of the retrograde transport, was obtained from Sigma, and used at 20 µM for 30 min as previously described ^55^. For antibody feeding, DLBCL cells were stained with anti-HSPG (10e4 reactivity) and subsequently with anti-mouse IgM TRITC at 4°C. Cells were incubated 30 min at 37°C in presence or in absence of recombinant APRIL. Cells were then treated for 3 min by an acid wash (Pierce^TM^ IgG elution buffer, pH 2.0) to remove surface staining. Finally, cells were incubated back at 37°C for 1 h and analyzed by confocal microscopy for HSPG expression. For cytokine quantification, 0.5 x 10^6^ DLBCL cells per ml were incubated for 48h for supernatant conditioning. Cell-free supernatants were analyzed for cytokine secretion using commercially available cytometric bead array using manufacturer’s instructions (BD biosciences). For mixed lymphocyte reactions, peripheral blood mononuclear cells (PBMC) from healthy donor (Etablissement Français du Sang, Grenoble) were isolated using standard Ficoll-paque centrifugation and stained with CellTrace CFSE using manufacturer’s instructions (Thermo Fisher). DLBCL cells were treated with APRIL and EDA-control for 2 days, and then incubated with CFSE-labeled PBMC at a 1:2 ratio. CD4^+^ T-cell division was evaluated by CFSE dilution 4 days later by flow cytometry.

### Cell transfection

Transient tranfection was performed in HEK 293-T cells (American Tissue Cell Collection) using the polyethylenimide (Sigma) method. BJAB and SU-DHL7 were transfected by electroporation with an electrosquareporator ECM 830 apparatus (BTX) with a single pulse of 250 V for 25 ms in 0,4 mm cuvettes. Constructs for Flag-tagged huAPRIL^A88^, flag-tagged huAPRIL^H98^, flag-tagged huEDA, full length huBCMA, GPI-anchored huBCMA and full length TACI were previously described ^35^. Construct encoding for full-length huBCMA fused to YPP was performed as previously described for other TNF receptors ^56^. Full-length murine BCMA (aa 1-184) was cloned in the PCR3 plasmid (Invitrogen). The IkBα mutant with a flag tag at its n-terminus encoded by the pMSCV vector was previously described ^57^. EGFP was subcloned in this vector by standard DNA subcloning methods. Stable BJAB cell transfectants was generated after electroporation as described above followed by selection with 1 µg/ml of puromycin (Invivogen). Transfected lines were cloned by limiting dilution and flag-reactivity among clones was assessed by Western blot analysis with biotinylated M2 anti-flag antibody (10 µg/ml) and HRP-conjugated streptavidin.

### Flow cytometry

Total staining was performed by using the transcription factor buffer kit using manufacturer’s instructions (BD biosciences). For internalization studies, cells were first incubated with recombinant flag-tagged huAPRIL^A88^ at 4°C for 45 min and then split in two, one part remaining at 4°C and the other one incubated at 37°C for the indicated time. Detection was performed in two steps with biotinylated anti flag (clone M2, Sigma) and fluorochrome-conjugated streptavidin (BD Biosciences). Data were acquired on a LSR II or Accurri C6plus (Becton Dickinson) and analyzed with FCS Express 6 (DeNovo software). Grey-shaded histograms correspond to isotype control stainings. A FACS-ARIA was used to purify cells.

### Cytofluorescence, immunohistofluorescence/chemistry and cytology

For cytofluorescence, DLBCL cells were incubated with recombinant APRIL at 4°C or 37°C and harvested in cold PBS. Cells were then fixed in 4% PFA and quenched with 0.1 M glycine in PBS. Cells were next permeabilized in PBS 2% BSA, 0.2% saponin and subsequently incubated with primary and secondary antibodies for 1 h at room temperature (RT) each. Hoechst 33342 (5 µg/ml, Thermo Fisher) was used to stain nuclei. Cells were mounted on a glass slide and analyzed by confocal microscopy on a Zeiss LSM 510 META with a plan-apochromat 63x/1.4 objective. Co-localization was analyzed by using “RG2B co-localization” plugin for ImageJ. Auto thresholding was used and pixel intensity acquired with measure tool for processed and original images. Ratio between the two measures determined co-localization percentage. Quantification was performed on at least 30 cells per condition and per experiment. Immunohistofluorescence/chemistry on formalin-fixed paraffin-embedded (FFPE) samples were performed as previously described ^18^. The rabbit polyclonal anti-BCMA was used after heat-induced epitope retrieval in citrate buffer. Cytoblocks with transfected cells were made after formalin fixation and paraffin embedment. Cytology of purified cells were performed on cytospins with May-Grünwald-Giemsa staining. Cells/tissues were visualized with an AxioImager M2 microscope (Zeiss).

### Immunoprecipitation and Western-blot

300 x 10^6^ BJAB cells were incubated with 50 µg of Flag-tagged APRIL for 25 min at 4°C. Cells were next incubated at 37°C in 6 ml serum-free OptiMEM medium, and 1 ml was harvested each 10 min and kept at 4°C until lysis with NP-40 lysis buffer (20 mM Tris-HCL, 150 mM NaCl, 10% glycerol, 1% NP-40, pH 7.5). Lysis buffer was supplemented with a protease inhibitor cocktail (cOmplete^TM^ ULTRA tablets, Roche) and Pierce^TM^ phosphatase inhibitor (Thermo Fisher). APRIL was precipitated overnight at 4°C with anti-Flag (clone M2) agarose beads (Sigma) and after extended washes, eluted in loading buffer (187.5 mM Tris-HCL, 6% SDS, 0.03% phenol red, 10% glycerol, pH 6.8) at 95°C for 10 min. Lysate before immunoprecipitation was used as input control and for NF-κB activation analysis. Migration was performed on a SDS polyacrylamide gel and proteins transferred to a PVDF membrane (Bio-Rad). Primary antibodies were incubated overnight at 4°C. Secondary antibodies conjugated to HRP (Santa Cruz technology) were incubated 1 h at RT. Revelation was performed with the clarity western ECL substrate (Bio-Rad), and chemiluminescent signal was acquired with a ChemiDoc^TM^ (Bio-Rad).

### RT-qPCR

Total RNA was extracted from cells with the RNeasy micro kit (Qiagen). cDNA was generated using oligo-dT_12-18_ and SuperScript II reverse transcriptase (Thermo Fisher). Quantification was performed using the BioRad CFX96 system and iTaq universal SybrGreen Supermix (Bio-Rad). The intron spanning forward 5’-atgggtcaggtggtgtctcg-3’, reverse 5’-tccccttggtgtaaatggaaga-3’ primers and the forward 5’-gctgtcatcgatttcttccc-3’, reverse 5’-ctcatggctttgtagatgcct-3’ primers were used for the detection of human *APRIL* and human *IL-10*, respectively. Expression levels were normalized using *actin* mRNA using forward 5’-agagggaaatcgtgcgtga-3’ and reverse 5’-gctggaaggtggacgtgag-3’primers. Results were quantified using a standard curve generated with serial dilutions of input DNA.

### Interaction of mouse APRIL with human BCMA

A standard ELISA was performed by coating human BCMA-Fc (10 µg/ml, Enzo Life Sciences) to test the interaction of flag-tagged-mouse/human APRIL^A88^. Cytotoxicity on JKT-huBCMA/Fas was performed as previously described ^28^

### Gene expression profiling

GEP for BCMA mRNA in healthy B cells was performed according to Genomicscape as described ^58^.

### Human and mouse experimentations

The Grenoble ethics committee approved mouse experimentation. Mouse experimentation was performed according to the ARRIVE guidelines. 0.5 x 10^6^ DLBCL cells were injected subcutaneously in 0.1 ml PBS. Tumor size was calculated using a caliper. When size was above 0,5 cm2, mice were euthanized. The generation of *APRIL*^-/-^ NOD/SCID mice have already been described ^9^. Female mice between 6 to 9 months were used. No animal randomization was performed. Tumor growth was monitored in blinded manner. Experimentation with primary fresh and FFPE human DLBCL samples was approved by the Grenoble-Alpes ethics committee and conducted according to the declaration of Helsinki. Samples were obtained after patients’ informed consent. Tonsils samples were obtained at the ORL department of the Grenoble-Alpes and Geneva University Hospitals from patients with acute tonsillitis and recurrent tonsillitis associated to infections and from patients with snoring problems from 2008 to 2015. For the latter, the patients had not experienced any tonsil infections for the previous 2 years and had not been treated with antibiotics during the last 3 months preceding surgery.

### Statistics

Statistical analyses were performed with t-tests with two group experiments and one-way Anova test with experiments containing superior than two groups using the Prism software (Graphpad Software Inc., San Diego, CA, USA). Geisser-Greenhouse corrections were performed in Anova tests. Welch corrections were performed in unpaired *t* tests. Normality was assessed with Shapiro-Wik tests. Parametric and non-parametric tests were performed with normally and abnormally distributed data, respectively. Specific tests are defined in figure legends. *P* values superior to 0.05 were considered as non-significant. * *p*<0,05, ** *p*<0,01, *** *p*<0,0001, **** *p*<0,00001.

