## supplemental figures for "BCMA signaling from the Golgi apparatus in B cells"

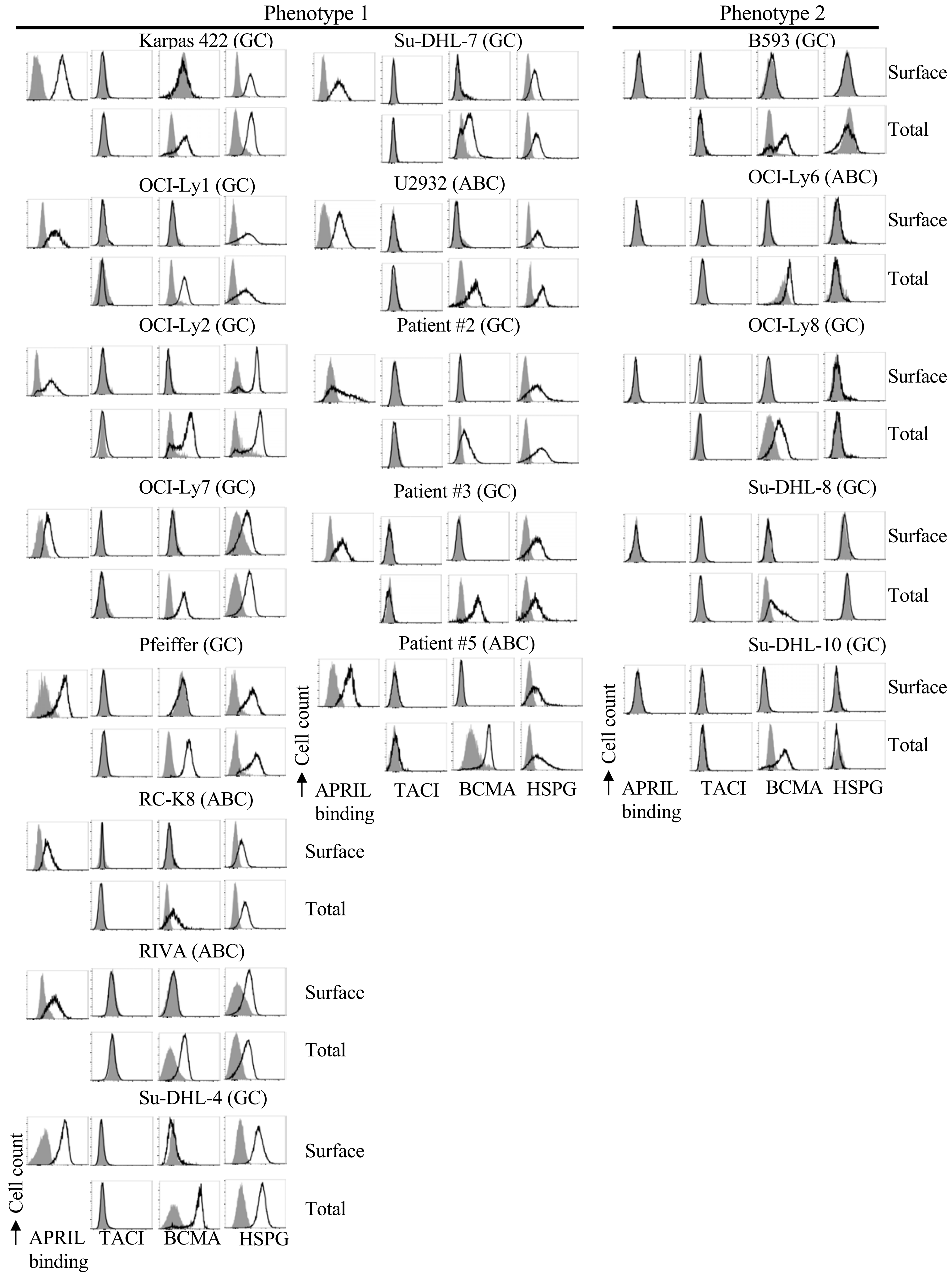
Phenotype 1
Phenotype 2
Karpas 422 (GC)
Su-DHL-7 (GC)
B593 (GC)
OCI-Ly1 (GC)
U2932 (ABC)
OCI-Ly6 (ABC)
OCI-Ly2 (GC)
Patient #2 (GC)
OCI-Ly8 (GC)
OCI-Ly7 (GC)
Patient #3 (GC)
Su-DHL-8 (GC)
Pfeiffer (GC)
Patient #5 (ABC)
Su-DHL-10 (GC)
RC-K8 (ABC)
RIVA (ABC)
Su-DHL-4 (GC)
Surface
Total
Cell count
APRIL binding
TACI
BCMA
HSPG

**Figure 1sup: GC and ABC DLBCL binds APRIL and harbors intracellular BCMA and surface HSPGs.**

All cell lines and patient were stained with recombinant $APRIL^{A88}$ and analyzed for the surface and total expression of the indicated receptors by flow cytometry. All **r**esults are representative of two independent experiments.

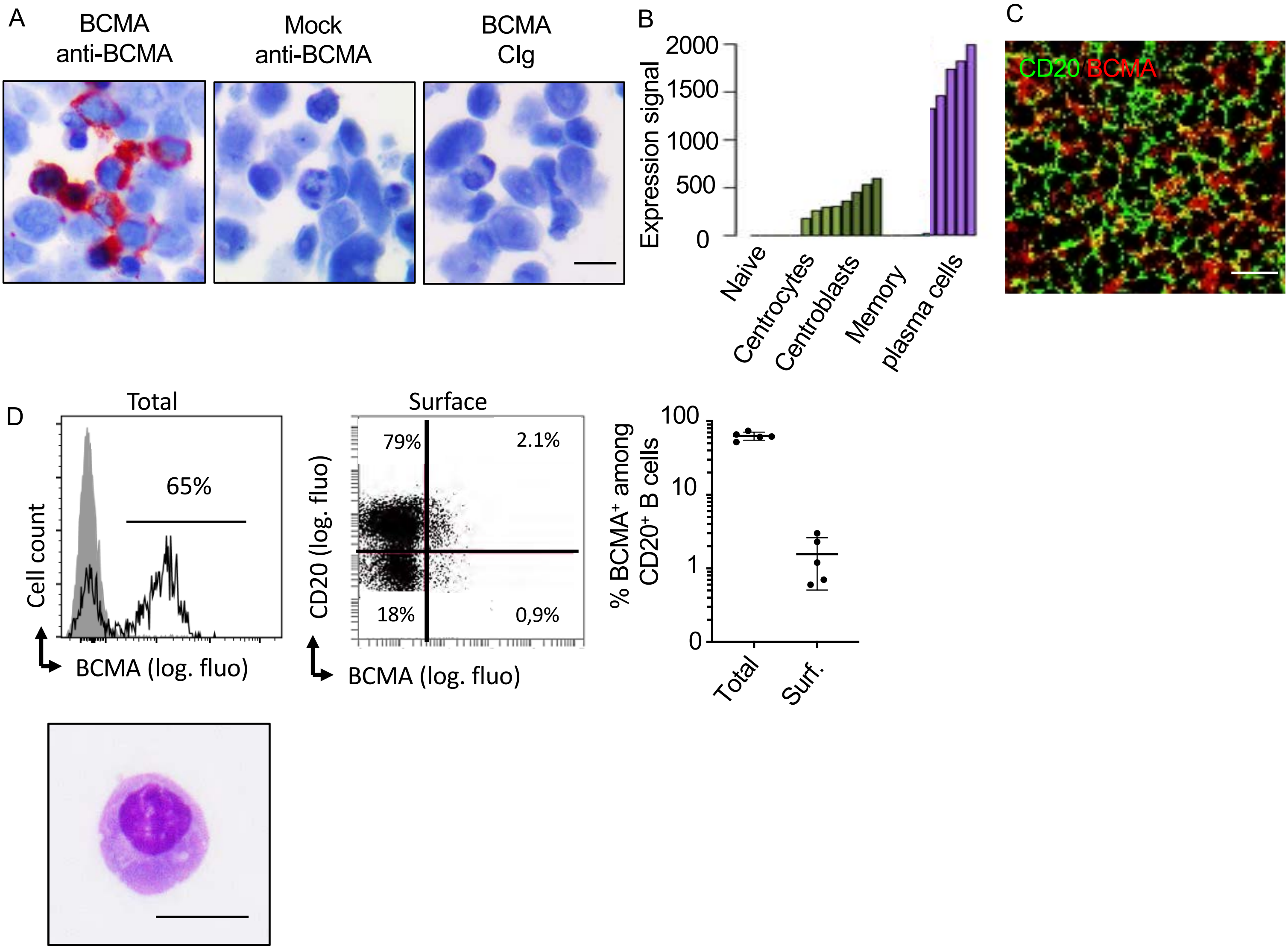


**Figure 2sup: Most GC B cells express intracellular BCMA**

A) Mock- and BCMA-transfected 293T cells were formalin-fixed and embedded into paraffin. Sections were stained by immunohistochemistry with a control Ig (cIg) and the anti-BCMA. Scale bar: 5 µm. Results are representative of two independent experiments. B) Gene expression profile is shown for BCMA (probe 206641_at) on purified naïve ($CD19^+CD27^-$) centrocytes ($CD19^+IgD^-CD38^+CXCR4^-$) and centroblasts ($CD19^+IgD^-CD38^+CXCR4^-$) from tonsils. Memory B cells ($CD19^+CD27^+$) and plasma cells ($CD138^+$) were from blood and bone, respectively. C) Tonsil sections were costained for CD20 and BCMA and analyzed by confocal microscopy. A germinal center zone is shown. Scale bar: 10 µm. D) Permeabilized and viable cells from tonsils were stained for CD20 and BCMA and analyzed by flow cytometry. Representative overlaid histogram plot from gated $CD20^+$ cells and dot plot are shown for total and surface stainings, respectively (upper left and middle panels). % of cells positive for BCMA is shown from five tonsils (upper right panel). Cytology of FACS-sorted surface $BCMA^+CD20^+$ cells from tonsils was analyzed by May-Grünwald-Giemsa staining (bottom panel). Scale bar: 5 µm. Results are representative of 12 different tonsils. FACS-sorting of surface $BCMA^+$ B cells was performed on 2 tonsils.

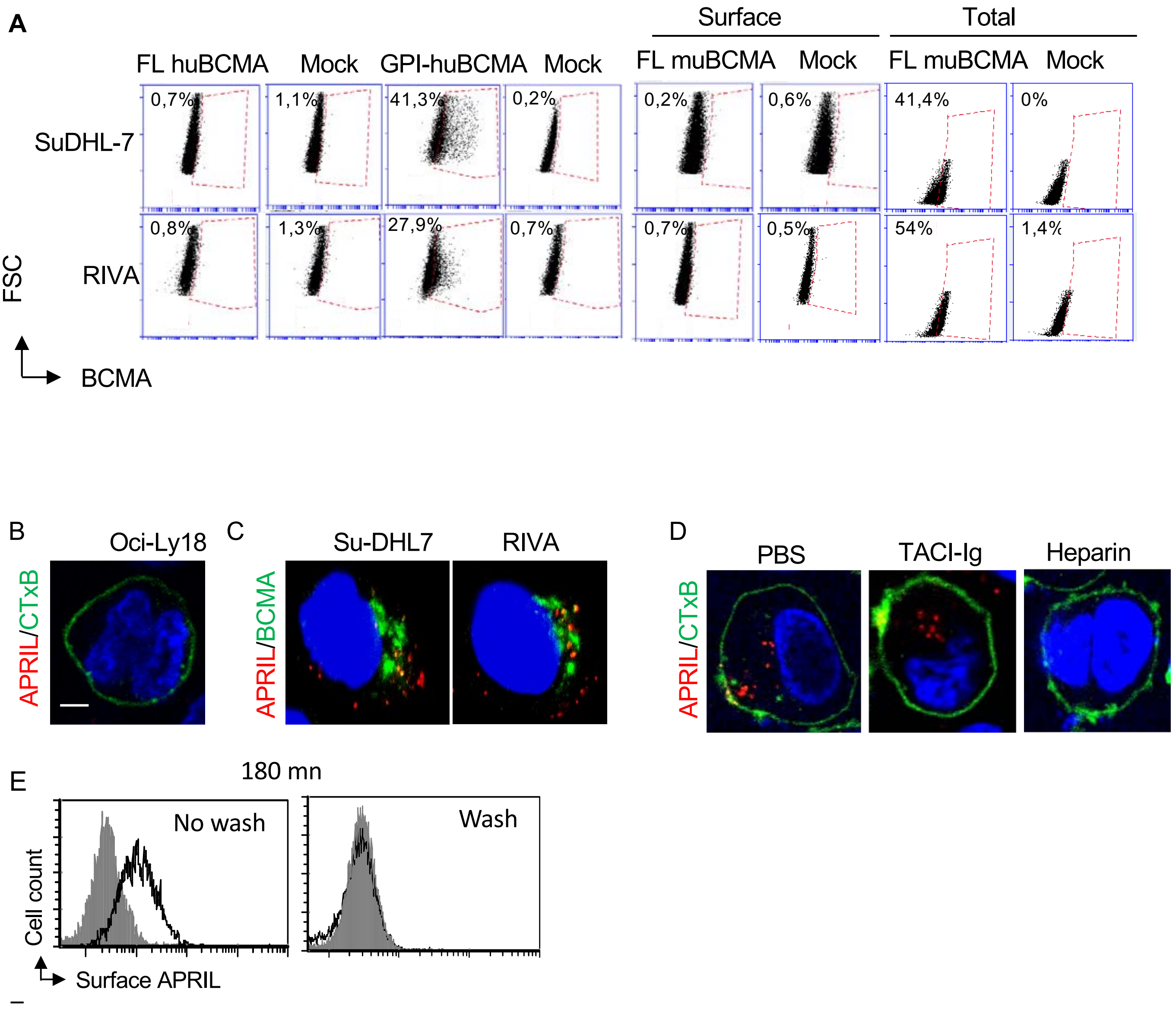


**Figure 3sup: BCMA is retained in the Golgi apparatus and colocalized with internalized APRIL in the Golgi apparatus.**

A) Cells transfected with the indicated constructs were analyzed after 48 hr by flow cytometry after surface or total staining of human and mouse BCMA. Dot plots are shown. FL: full length, GPI: glycophosphatidyl inositol. B) $APRIL^{A88}$ was added to Oci-LY18 from phenotype 2 at 4°C. Cells were washed and incubated at 37°C for 15 mn. Anti-flag reactivity (red) and CTxB localization (green) were assessed after cell permeabilization. C) $APRIL^{A88}$ was added to Su-DHL7 and RIVA from phenotype 1 at 4°C. Cells were washed and incubated at 37°C for 45 mn. Anti-flag (red) and anti-BCMA (green) reactivities were assessed after cell permeabilization. D) $APRIL^{A88}$ pretreated with the indicated reagents was added to BJAB cells at 4°C. Cells were washed and incubated at 37°C for 45 mn. Anti-flag reactivity (red) and CTxB (green) was assessed after cell permeabilization. Scale bar = 2µm. E) $APRIL^{A88}$ was added to BJAB cells at 4°C. Cells were washed or not and incubated at 37°C for 180 mn prior to staining for surface APRIL and flow cytometry analysisResults in A/B are representative of two independent experiments. Results in A/B and C/D/E are representative of two and three, respectively, independent experiments.

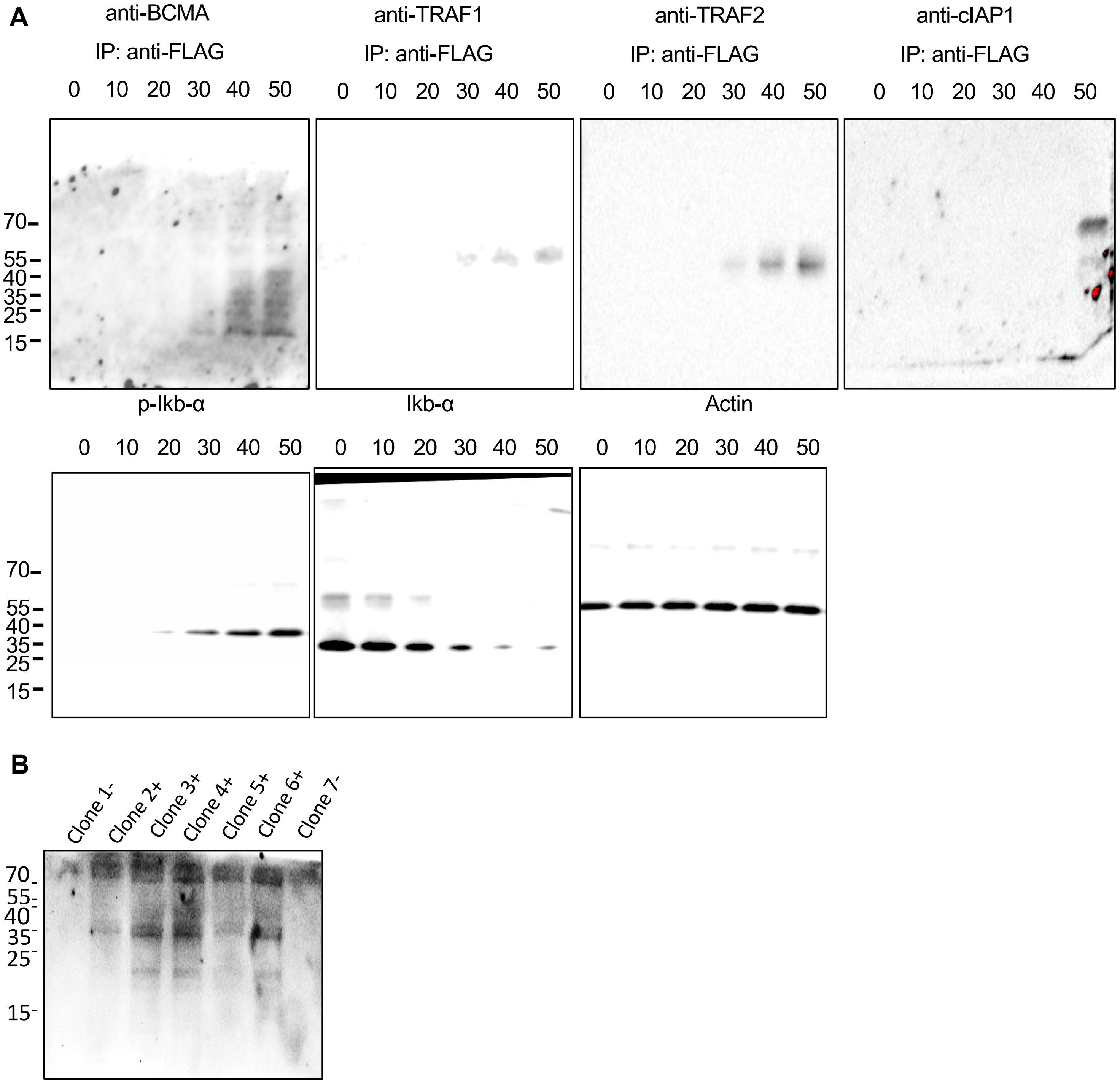


**Figure 4sup: uncropped Western blots**

A) For figure 6A. B) For figure 6D. Positivity for the IkB mutant is indicated.

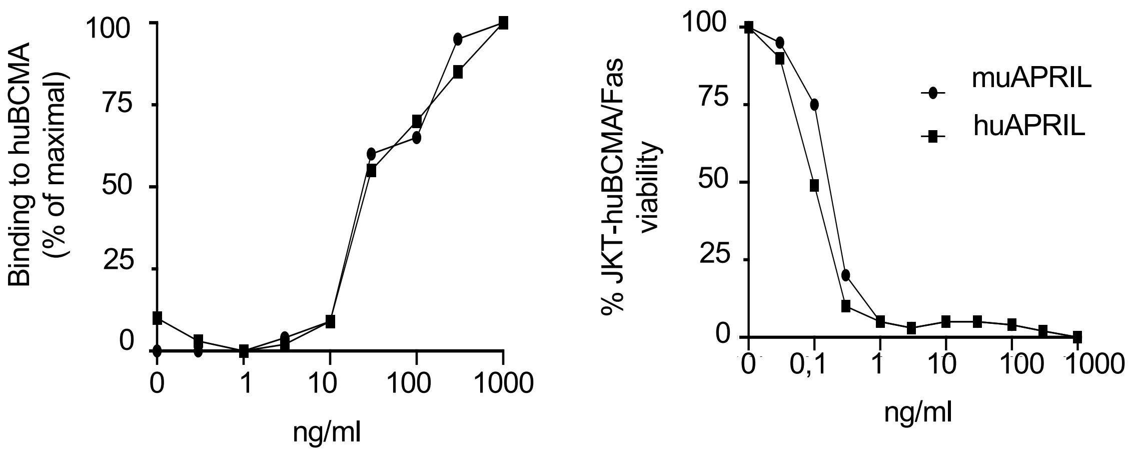


**Figure 5sup: MuAPRIL binds and stimulates huBCMA**

Binding of the indicated concentrations of hu and mu flag-ACRP-APRIL$^{A88}$ onto coated huBCMA was assessed in ELISA (left panel). % of max was defined by the OD value obtained with the highest concentration of huAPRIL used. Induction of cell death in jurkat (JKT) cells transfected with the reporter huBCMA/FAS by the indicated concentrations of hu and mu APRIL.
